# Antibody Neutralization of Emerging SARS-CoV-2: EG.5.1 and XBC.1.6

**DOI:** 10.1101/2023.08.21.553968

**Authors:** Qian Wang, Yicheng Guo, Richard M. Zhang, Jerren Ho, Hiroshi Mohri, Riccardo Valdez, David M. Manthei, Aubree Gordon, Lihong Liu, David D. Ho

## Abstract

SARS-CoV-2 variants EG.5.1 and XBC.1.6 have recently emerged, attracting increased attention due to their rapid expansion globally and in Australia, respectively. EG.5.1 evolved from Omicron subvariant XBB.1.9, harboring additional Q52H and F456L spike substitutions. The F456L mutation is located within the epitopes of many class-1 monoclonal antibodies (mAbs) directed to the receptor-binding domain (RBD), raising concerns about further antibody evasion. XBC.1.6, a descendant of a Delta-BA.2 recombinant, carries 15 additional spike mutations. The extent to which antibody evasion contributes to the growth advantage of XBC.1.6 in Australia remains to be determined. To assess the antibody evasion properties of the emergent variants, we conducted pseudovirus neutralization assays using sera from individuals who received three doses of COVID-19 mRNA monovalent vaccines plus one dose of a BA.5 bivalent vaccine, as well as from patients with BQ or XBB breakthrough infection. The assays were also performed using a panel of 14 mAbs that retained neutralizing activity against prior XBB subvariants. Our data suggested that EG.5.1 was slightly but significantly more resistant (< 2-fold) to neutralization by BQ and XBB breakthrough sera than XBB.1.16, which is known to be antigenically similar to XBB.1.5. Moreover, the F456L mutation in EG.5.1 conferred heightened resistance to certain RBD class-1 mAbs. In contrast, XBC.1.6 was more sensitive to neutralization by sera and mAbs than the XBB subvariants. Notably, XBB breakthrough sera retained only weak neutralization activity against XBB subvariants. In summary, EG.5.1 and XBC.1.6 exhibited distinct antibody evasion properties. The recent global expansion of EG.5.1 might be attributable, in part, to its enhanced neutralization resistance. That XBB breakthrough infections did not elicit a robust antibody neutralization response against XBB subvariants is indicative of immunological imprinting. The high prevalence of XBC.1.6 in Australia is not due to enhanced antibody evasion.

## Main text

Although the World Health Organization has declared an end to the emergency phase of the COVID-19 pandemic, SARS-CoV-2 has continued to spread and evolve, giving rise to novel variants^1^. These new forms contained additional spike mutations that could further compromise protection mediated by antibodies elicited by prior infections and/or vaccinations. Recently, two emergent Omicron subvariants, EG.5 and EG.5.1, have expanded rapidly worldwide, including in the United States but particularly in China **(Figure 1A)**. EG.5 evolved from Omicron XBB.1.9 (**Figure S1A)** and harbors one additional F456L substitution in the receptor-binding domain (RBD) of spike relative to the recently dominant subvariant XBB.1.5 (**Figure S1B)**. Its immediate descendant, EG.5.1, contains one more mutation, Q52H in the N-terminal domain (NTD) of spike **(Figure S1B)**. Notably, both new subvariants possess the F456L mutation, which could potentially evade some of the antibodies targeting the so-called class-1 region of RBD^2^. Another emergent subvariant is XBC.1.6, a recombinant between Delta and Omicron BA.2 (**Figure S1A)** that has gained prevalence in Australia (**Figure 1A)**. It has 9 NTD mutations specific to the Delta variant as well as 6 more mutations in comparison to the BA.2 spike **(Figure S1C)**. The local growth advantage of XBC.1.6 underscores the need to understand its antibody evasion properties.

**Figure 1.**
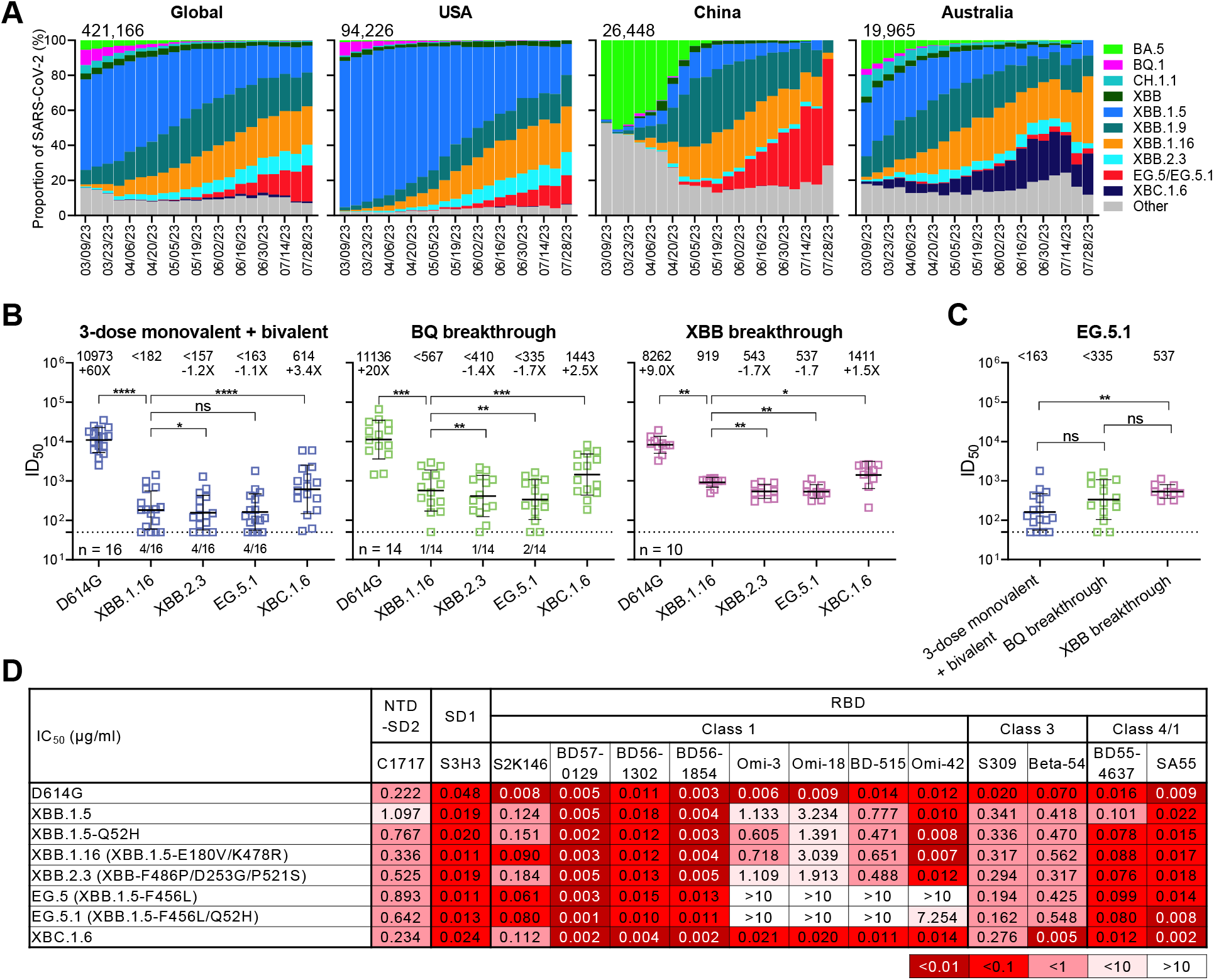
Prevalence of SARS-CoV-2 variants and their resistance to neutralization by human sera and monoclonal antibodies. **A.** Frequencies of currently circulating SARS-CoV-2 variants as of August 9, 2023. The number in the upper left corner of each graph reflects the cumulative viral sequences deposited to GISAID during the specified period, either globally or within the indicated countries. **B**. Serum neutralizing ID_50_ titers from “3-dose monovalent + Bivalent”, “BQ breakthrough” and “XBB breakthrough” cohorts. Values above the symbols denote the geometric mean ID_50_ titer (GMT) for each cohort. Comparisons show the fold change in GMT relative to that of XBB.1.16. Statistical significance was analyzed by Wilcoxon matched-pairs signed-rank tests. The limit of detection (LOD) of the assay is 50, as shown by a dotted line. The number of samples at or below the LOD is denoted above the X-axis, and n represents sample size. **C**. Serum neutralizing ID_50_ titers against EG.5.1 are displayed for three indicated cohorts. Statistical analyses were performed using Mann-Whitney unpaired tests. **D**. Pseudovirus neutralization IC_50_ values for monoclonal antibodies against indicated variants and one point mutant in comparison with D614G and XBB.1.5. Significance levels are annotated as: ns, not significant; *, *p* < 0.05; **, *p* < 0.01; ***, *p* < 0.001 and ****, *p* < 0.0001.

We evaluated the neutralization of EG.5.1 and XBC.1.6 by sera from three different clinical cohorts (**Table S1)**. The ancestral virus D614G and subvariants XBB.2.3 and XBB.1.16 were included as comparators. The first cohort comprised of individuals who received one of the BA.5 bivalent COVID-19 mRNA vaccines after receiving three doses of one of the original COVID-19 mRNA vaccines (termed “3 shots WT + bivalent”). The other two cohorts included individuals who experienced a BQ or XBB breakthrough infection after multiple vaccinations (termed “BQ breakthrough” or “XBB breakthrough”, respectively). In all three groups, the XBB subvariants (XBB.1.16, XBB.2.3, and EG.5.1) were substantially more resistant to serum neutralization than D614G, while XBC.1.6 showed greater sensitivity to serum neutralization compared to the XBB subvariants **(Figure 1B)**. In particular, EG.5.1 displayed a small (1.7-fold) but significant increase in resistance to serum neutralization for the breakthrough cohorts when compared with XBB.1.16, which is known to have similar antibody neutralization profile as XBB.1.5^3^. These serum neutralization results were then used to generate antigenic maps to reflect the antigenic relationships among the SARS-CoV-2 variants evaluated (**Figure S2**). Moreover, serum neutralization titers against EG.5.1 were modestly higher for the XBB breakthrough subjects but relatively low overall in all group **(Figure 1C)**, perhaps due to persistence of immunological imprinting^3,4^.

We next tested antibody evasion properties of EG.5, EG.5.1, and XBC.1.6 using a panel of monoclonal antibodies that retained neutralizing activity against XBB.1.5 by targeting a number of epitope clusters on spike. D614G, XBB.1.5, XBB.1.16, XBB.2.3, and XBB.1.5 with Q52H point mutation were included as comparators. XBB.1.5, XBB.1.16 and XBB.2.3 exhibited similar neutralization profiles, while EG.5 and EG.5.1 showed marked neutralization resistance to 4 of 8 RBD class-1 monoclonal antibodies (**Figure 1D**), undoubtedly mediated by the F456L mutation. Structural modeling revealed that this mutation reduced the size of the side chain and thus decreased the hydrophobic interaction with CDRH3 of antibodies Omi-3, Omi-18, and BD-515 (**Figure S3A-C**). Furthermore, the F456L mutation resulted in a clash with residue F101 in the CDRH3 of Omi-42, leading to loss of neutralization (**Figure S3D**). Remarkably, as compared to XBB subvariants, XBC.1.6 was more sensitive to neutralization by half of the RBD monoclonal antibodies (**Figure 1D**), consistent with results from serum neutralization. Point mutation Q52H (found in EG.5.1 but not EG.5) did not have an appreciable impact on antibody neutralization when introduced into the background of XBB.1.5, a finding in agreement with similar neutralization profiles for EG.5 and EG.5.1 (**Figure 1D**).

In summary, our findings indicate that the recently surging subvariants EG.5 and EG.5.1 are only modestly (1.7-fold) more resistant to neutralization by serum antibodies than the previously dominant subvariant XBB.1.5, largely due to the F456L mutation in the viral spike knocking out the binding of some of the antibodies that target the class-1 region of RBD. It is not likely that these new subvariants will have a dramatically adverse impact on the efficacy of current COVID-19 vaccines compared to their predecessor XBB.1.5. Although an extra degree of antibody resistance may confer a growth advantage to EG.5/EG.5.1, it is also possible that mutations elsewhere in the genome are contributing, such as the I5T mutation in Orf9b^5^. On the other hand, XBC.1.6 is appreciably more sensitive to antibody neutralization than XBB subvariants; therefore, it is less likely to compromise vaccine efficacy. Nevertheless, it is puzzling why XBC.1.6 is gaining in prevalence in Australia over XBB subvariants that are noticeably more evasive to antibodies. As SARS-CoV-2 continues to spread and mutate, it is imperative that we remain vigilant in tracking its evolutionary trajectory as well as in understanding the functional consequences of its mutations.

