## Supplemental information for "Antibody Neutralization of Emerging SARS-CoV-2: EG.5.1 and XBC.1.6"

#### Supplementary Appendix

### Supplementary Appendix

#### Table S1. Demographics of clinical cohorts.

| Sample ID | Vaccine type and infected strain | Days post-bivalent vaccination or *infection | Documented COVID-19 | Age | Gender |
|---|---|---|---|---|---|
| 3 shots WT + bivalent | | | | | |
| UM-36 | BNT162b2/BNT162b2/BNT162b2/Moderna Bivalent | 24 | No | 38 | Female |
| UM-37 | BNT162b2/BNT162b2/BNT162b2/Moderna Bivalent | 27 | No | 42 | Female |
| UM-39 | mRNA-1273//mRNA-1273/mRNA-1273/Moderna Bivalent | 24 | No | 36 | Male |
| UM-43 | BNT162b2/BNT162b2/BNT162b2/Pfizer Bivalent | 25 | No | 49 | Female |
| UM-44 | BNT162b2/BNT162b2/BNT162b2/Moderna Bivalent | 25 | No | 37 | Female |
| UM-47 | BNT162b2/BNT162b2/BNT162b2/Pfizer Bivalent | 26 | No | 45 | Male |
| UM-48 | BNT162b2/BNT162b2/mRNA-1273/Moderna Bivalent | 26 | No | 43 | Female |
| UM-51 | mRNA-1273/mRNA-1273/mRNA-1273/Moderna Bivalent | 29 | No | 32 | Female |
| UM-52 | BNT162b2/BNT162b2/BNT162b2/Pfizer Bivalent | 23 | No | 43 | Female |
| UM-56 | BNT162b2/BNT162b2/mRNA-1273/Moderna Bivalent | 27 | No | 36 | Female |
| UM-60 | BNT162b2/BNT162b2/BNT162b2/Moderna Bivalent | 30 | No | 24 | Female |
| Q101 | mRNA-1273/mRNA-1273/mRNA-1273/Moderna Bivalent | 30 | No | 32 | Female |
| Q102 | BNT162b2/BNT162b2/mRNA-1273/Moderna Bivalent | 23 | No | 39 | Male |
| Q103 | BNT162b2/BNT162b2/BNT162b2/Pfizer Bivalent | 30 | No | 26 | Female |
| Q104 | mRNA-1273/mRNA-1273/mRNA-1273/Pfizer Bivalent | 30 | No | 27 | Female |
| Q105 | BNT162b2/BNT162b2/BNT162b2/Pfizer Bivalent | 23 | No | 23 | Male |
| BQ breakthrough | | | | | |
| BQ-1 | BNT162b2/BNT162b2/BNT162b2/BNT162b2/mRNA-1273/BQ | *33 | Yes | 53 | Female |
| BQ-2 | BNT162b2/BNT162b2/BNT162b2/mRNA-1273/BQ | *30 | Yes | 32 | Female |
| BQ-3 | BNT162b2/BNT162b2/BNT162b2/mRNA-1273/BQ | *21 | Yes | 35 | Female |
| BQ-4 | BNT162b2/BNT162b2/mRNA-1273/BQ | *39 | Yes | 52 | Female |
| BQ-5 | BNT162b2/BNT162b2/BNT162b2/BNT162b2/mRNA-1273/BQ | *22 | Yes | 62 | Female |
| BQ-6 | BNT162b2/BNT162b2/mRNA-1273/BQ | *34 | Yes | 29 | Female |
| BQ-7 | BNT162b2/BNT162b2/BQ | *44 | Yes | 35 | Female |
| BQ-8 | mRNA-1273/mRNA-1273/mRNA-1273/mRNA-1273/BQ | *59 | Yes | 33 | Female |
| BQ-9 | BNT162b2/BNT162b2/BNT162b2/BNT162b2/BQ | *44 | Yes | 45 | Female |
| BQ-10 | JNJ-78436735/JNJ-78436735/BNT162b2/BNT162b2/BQ | *32 | Yes | 47 | Female |
| BQ-11 | BNT162b2/BNT162b2/BNT162b2/BQ | *36 | Yes | 36 | Female |
| BQ-12 | BNT162b2/BNT162b2/BNT162b2/BNT162b2/BNT162b2/BQ | *77 | Yes | 59 | Female |
| BQ-13 | BNT162b2/BNT162b2/BNT162b2/BNT162b2/BQ | *25 | Yes | 39 | Female |
| BQ-14 | BNT162b2/BNT162b2/BNT162b2/BNT162b2/BQ | *67 | Yes | 43 | Female |
| XBB breakthrough | | | | | |
| XBB-1 | BNT162b2/BNT162b2/BNT162b2/BNT162b2/BNT162b2/XBB | *35 | Yes | 38 | Female |
| XBB-2 | mRNA-1273/mRNA-1273/mRNA-1273/BNT162b2/Moderna bivalent/XBB | *50 | Yes | 38 | Male |
| XBB-3 | BNT162b2/BNT162b2/BNT162b2/Moderna bivalent/XBB | *54 | Yes | 61 | Male |
| XBB-4 | BNT162b2/BNT162b2/mRNA-1273/BNT162b2/XBB | *23 | Yes | 41 | Male |
| XBB-5 | BNT162b2/BNT162b2/BNT162b2/mRNA-1273/XBB | *14 | Yes | 37 | Male |
| XBB-6 | BNT162b2/BNT162b2/mRNA-1273/BNT162b2/XBB | *78 | Yes | 41 | Male |
| XBB-7 | BNT162b2/BNT162b2/BNT162b2/mRNA-1273/XBB | *79 | Yes | 37 | Male |
| XBB-8 | BNT162b2/BNT162b2/BNT162b2/BNT162b2/BNT162b2/XBB | *86 | Yes | 61 | Female |
| XBB-9 | mRNA-1273/mRNA-1273/mRNA-1273/BNT162b2/mRNA-1273/XBB | *103 | Yes | 41 | Female |
| XBB-10 | BNT162b2/BNT162b2/BNT162b2/mRNA-1273/XBB | *107 | Yes | 32 | Male |

**A**

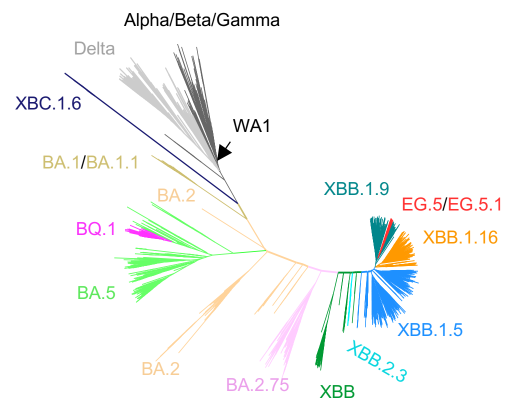

**B**

| VOCs | Q52 | E180 | G252 | D253 | F456 | K478 | S486 | P521 |
|---|---|---|---|---|---|---|---|---|
| XBB | | | | | | | | |
| XBB.1.5 | | | V | | | | P | |
| XBB.1.16 | | V | V | | | R | P | |
| XBB.2.3 | | | | G | | | P | S |
| EG.5 | | | V | | L | | P | |
| EG.5.1 | H | | V | | L | | P | |

**C**

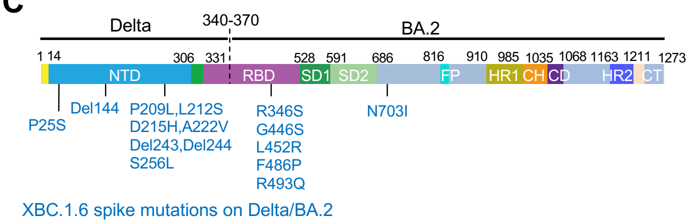

**Figure S1. Unrooted phylogenetic time tree of SARS-CoV-2 and spike mutations found in XBB subvariants and XBC.1.6.**

**A.** Unrooted phylogenetic time tree showing XBB subvariants and XBC.1.6 in relation to other main SARS-CoV-2 variants. The tree was download from https://nextstrain.org/ncov/gisaid/global/6m?l=unrooted.

**B.** Spike mutations found in XBB subvariants relative to XBB.

**C.** Spike mutations found in XBC.1.6 compared to Delta or BA.2. The dashed line marks the possible recombination breakpoint between Delta and BA.2 in XBC variant.

NTD, N-terminal domain; RBD, receptor-binding domain; SD1 and SD2, subdomains 1 and 2; FP, fusion peptide; HR1 and HR2, heptad repeat region 1 and 2; CH, central helical region; CD, connector domain; CT, cytoplasmic tail.

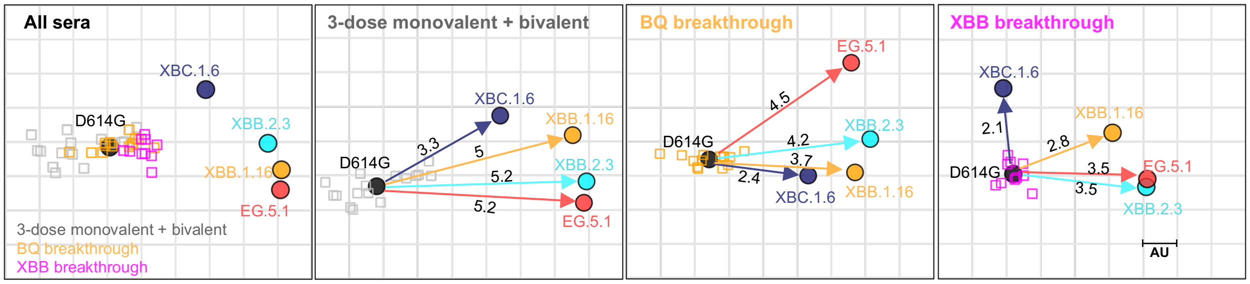

**Figure S2. Antigenic map generated by neutralization data from various serum cohorts as indicated.** D614G serves as the central reference for all serum cohorts, with the antigenic distances calculated to denote the average divergence from each variant. One antigenic unit (AU) represents an approximately 2-fold change in $ID_{50}$ titer.

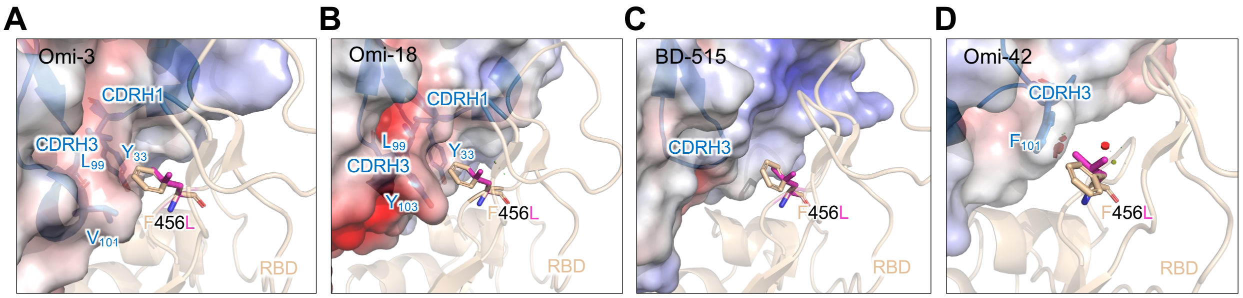

**Figure S3. Structural modeling of the impact of F456L on RBD class 1 mAb binding.**
Semi-transparent surfaces indicate the electrostatic potential of each antibody, the red plates highlight steric hindrance between the mutated RBD and the respective antibody.

**Supplementary Methods**

*Patients and vaccinees*

The serum samples in this study were obtained from three cohorts: the "3-dose monovalent + bivalent", "BQ breakthrough" and "XBB breakthrough". The first cohort were individuals who received three doses of monovalent COVID-19 mRNA vaccine (mRNA-1273 or BNT162b2) followed by one dose of the Pfizer or Moderna bivalent mRNA vaccine. The last two cohorts comprised of participants who had Omicron BQ or XBB breakthrough infection after vaccination. The "3-dose monovalent + bivalent" cohort were confirmed uninfected by history and anti- nucleocapsid protein (NP) ELISA. The sequencing was conducted to determine the viral genotype for "BQ breakthrough" and "XBB breakthrough" cohorts.

Serum samples were collected either at Columbia University Irving Medical Center or from the Immunity-Associated with SARS-CoV-2 Study (IASO), an ongoing cohort study that began in 2020 at the University of Michigan. All participants provided written informed consent, and the serum collections were performed under protocols reviewed and approved by the Institutional Review Board of Columbia University or the University of Michigan. Detailed clinical information for each donor is provided in Table S1.

*Cell lines*

HEK293T cells (CRL-3216) and Vero-E6 cells (CRL-1586) were purchased from the American Type Culture Collection (ATCC) and maintained in Dulbecco modified Eagle medium (DMEM) with 10% fetal bovine serum (FBS) and 1% penicillin-streptomycin. These two cell lines were cultured in an atmosphere of 5% $CO_2$ at 37°C.

Expi293 cells were purchased from Thermo Fisher Scientific (A14527) and cultured in Expi293™ Expression Medium supplemented with 0.5% penicillin-streptomycin at 37°C, 8% $CO_2$, and 125 rpm.

*Monoclonal antibodies*

Antibodies (C1717[1], S3H3[2], S2K146[3], BD57-0129[4], BD56-1302[4], BD56-1854[4], Omi-3[5], Omi-18[5], BD-515[6], Omi-42[5], sotrovimab[7], Beta-54[8], BD55-4637[4], SA55[9]) were expressed in-house as previously described[10]. Briefly, antibody heavy chain and light chain variable genes were synthesized from GenScript, transfected into Expi293 cells using polyethylenimine (PEI), and then purified from supernatants using rProtein A Sepharose (GE) 4 days post transfection.

*Construction of SARS-CoV-2 spike plasmids*

Plasmid constructs encoding D614G, XBB.1.5, and XBB.1.16 spikes were produced as previously described[11]. Using the QuikChange II XL site-directed mutagenesis kit from Agilent, constructs for XBB.2.3, EG.5, EG.5.1, XBB.1.5-Q52H, and XBB.1.5-N703I spikes were generated as per manufacturer guidelines. XBC.1.6 spike was procured via synthesis from GenScript. All constructs were confirmed by sequencing before use.

*Pseudovirus production*

Pseudotyped SARS-CoV-2 were generated as previously described. HEK293T cells were first transfected with a plasmid encoding the spike protein of choice using 1 mg/mL of PEI and cultured under 5% $CO_2$ at 37 °C for 24 hours. Then transfected HEK293T cells were infected with VSV-G pseudotyped ΔG-luciferase

(G*ΔG-luciferase, Kerafast) a multiplicity (MOI) of ~3 to 5. Two hours later, cells were washed three times with PBS and cultured in fresh medium for another 24 hours. The supernatants were then collected, clarified by centrifugation, and aliquoted for storage at -80 °C until use.

*Pseudovirus neutralization assay*

Pseudoviruses underwent titration to standardize the viral input prior to each neutralization assay. Both antibodies and heat-inactivated sera were subjected to serial dilutions, starting from 10 μg/mL and a 1:100 dilution, respectively. These dilutions were carried out using a five-fold dilution factor in triplicate in 96-well plates. Following this, pseudoviruses were added and incubated with varying dilutions of either the antibody or serum at 37 °C for 1 hour. Control wells solely containing the pseudovirus were incorporated in all assay plates. Subsequently, Vero-E6 cells were seeded at a density of $4 \times 10^4$ cells/well, and the neutralization plates were maintained at 37 °C overnight. Upon completion, cellular lysis was conducted, and the resultant luciferase activity was quantified employing the Luciferase Assay System (Promega) in tandem with the SoftMax Pro v.7.0.2 software (Molecular Devices), as per the guidelines provided by the manufacturers. Both $ID_{50}$ and $IC_{50}$ values were calculated via nonlinear five-parameter dose-response curve fitting using GraphPad Prism v.9.2.

*Antigenic cartography*

Antigenic distances between sera to D614G and other SARS-CoV-2 variants were determined by integrating ID50 values of individual serum samples through an established antigenic cartography approach[12]. The visualization was crafted using the Racmacs package (v.1.1.4, https://acorg.github.io/Racmacs/) in R version 4.0.3. With optimization steps set at 2,000 and the minimum column basis parameter set to 'none', the 'mapDistances' function was employed to calculate antigenic distances between each serum sample and variant. Average distances from all sera to each variant determined the final representation. D614G served as the center of sera for each group, the seeds for each antigenic map were manually adjusted to ensure that EG.5.1 was displayed in the horizontal direction relative to the sera.

*Structural modeling*

The structures of antibody–RBD complexes for modelling were obtained from PDB (7ZF3 (Omi-3), 7ZFB (Omi-18), 7EE8 (BD515) and 7ZR7 (Omi-42)). The electrostatic potential was estimated by APBS electrostatics plugin, and the mutagenesis analysis were performed by Pymol version 2.5.4 (Schrödinger, LLC).

*Quantification and statistical analysis*

$ID_{50}$ values for serum neutralization and $IC_{50}$ values for antibody neutralization were obtained from a five-parameter dose-response curve using GraphPad Prism v.9.2. Statistical assessments were performed by either Wilcoxon matched-pairs signed-rank tests or Mann-Whitney unpaired tests in the same software. Significance levels are annotated as: ns, not significant; *, $p < 0.05$; **, $p < 0.01$; ***, $p < 0.001$ and ****, $p < 0.0001$.

**Acknowledgements**

This study was supported by funding from the NIH SARS-CoV-2 Assessment of Viral Evolution (SAVE) Program (Subcontract No. 0258-A709-4609 under Federal Contract No. 75N93021C00014) attributed to D. D. H., as well as the NIH, NIAID under contract number 75N93019C00051 attributed to A.G.. We thank the contributors for uploading sequences to the GISAID database.

**Author Contributions**

L.L. and D.D.H. conceived the study. Q.W., R.Z., J.H., J. Y., and L.L. performed experiments. H.M., R.V., E.S. and A.G. collected serum samples. Y.G. performed bioinformatics analysis. Q.W., Y.G., L.L., and D.D.H. analyzed the results and wrote the manuscript. L.L. and D.D.H. directed and supervised the project. All authors reviewed and approved of the manuscript.

**Declaration of Interests**

D.D.H. is a co-founder of TaiMed Biologics and RenBio, consultant to WuXi Biologics and Brii Biosciences, and board director for Vicarious Surgical. A.G. served on a scientific advisory board for Janssen Pharmaceuticals. The remaining authors declare no competing interests.
